# Cooperative Accumulation of Dynein-Dynactin at Microtubule Minus-Ends Drives Microtubule Network Reorganization

**DOI:** 10.1101/140392

**Authors:** Ruensern Tan, Peter J. Foster, Daniel J. Needleman, Richard J. McKenney

## Abstract

**Summary:** Cytoplasmic dynein-1 (dynein) is minus-end directed motor protein that transports cargo over long distances and organizes microtubules (MTs) during critical cellular processes such as mitotic spindle assembly. How dynein motor activity is harnessed for these diverse functions remains unknown. Here, we have uncovered a mechanism for how processive dynein-dynactin complexes drive MT-MT sliding, reorganization, and focusing, activities required for mitotic spindle assembly. We find that motors cooperatively accumulate, in limited numbers, at MT minus-ends. Minus-end accumulations drive MT-MT sliding, independent of MT orientation, and this activity always results in the clustering of MT minus-ends. At a mesoscale level, activated dynein-dynactin drives the formation and coalescence of MT asters. Macroscopically, dynein-dynactin activity leads to bulk contraction of millimeter-scale MT networks, demonstrating that minus-end accumulations produce network scale contractile stresses. Our data provides a model for how localized dynein activity is harnessed by cells to produce contractile stresses within the mitotic spindle.

**Highlights:**

- Processive dynein-dynactin complexes cooperatively form stable accumulations at MT minus-ends.
- Minus-end accumulations of motors slide MTs without orientation bias, leading to minus-end focusing.
- Minus-end accumulations of motors organize dynamic MTs into asters.
- Minus-end accumulations of motors drive bulk contractions of large-scale MT networks.

## Introduction

Cytoplasmic dynein-1 (dynein), is the only minus-end directed microtubule (MT) motor complex in animal cells capable of processive, long-distance transport (Allan, 2011; Cianfrocco et al., 2015; Vallee et al., 2012). The ~1.4 MDa, multi-subunit dynein complex is the largest and most complex of the MT motor proteins and belongs to the AAA+ family of ring-shaped molecular motors (Carter et al., 2016; Vale, 2000). It is responsible for a wide variety of cellular functions and transports many types of cargo, including membrane-bound organelles, mRNAs, stress granules, viruses, and misfolded proteins (Cianfrocco et al., 2015). In addition to canonical cargo transport, dynein has been implicated in intracellular MT organization. In neurons, dynein activity is necessary to maintain the correct polarity of axonal MTs (Zheng et al., 2008). In mitosis, dynein plays various roles including MT organization (Heald et al., 1996; Merdes et al., 2000; Rusan et al., 2002), transport of mitotic checkpoint proteins from kinetochores (Bader and Vaughan, 2010; Howell et al., 2001; Whyte et al., 2008), cortical pulling of astral MTs (McNally, 2013), tethering of centrosomes to spindle poles (Goshima et al., 2005; Maiato et al., 2004; Morales-Mulia and Scholey, 2005), and regulation of kinetochore MT attachment (Varma et al., 2008).

During self-organization of the mitotic spindle, dynein activity antagonizes kinesin-5-, and kinesin-12-dependent extensile MT-MT sliding (Ferenz et al., 2009; Mitchison et al., 2005; Tanenbaum et al., 2008; Tanenbaum et al., 2009; Vanneste et al., 2009). Indeed, inhibition of kinesin-5 results in dynein-driven spindle collapse into a monopolar structure (Ferenz et al., 2009; Tanenbaum et al., 2008). Dynein activity is also required to dynamically integrate and focus MT minus-ends into the spindle poles (Elting et al., 2014; Sikirzhytski et al., 2014), in both the presence and absence of centrosomes (Goshima et al., 2005; Heald et al., 1996). While dynein’s functions within the spindle are defined, the molecular mechanism for how it performs these activities remains to be fully understood. Dynein cooperates with nuclear mitotic apparatus protein 1 (NuMA) to slide MTs within the spindle (Merdes et al., 2000; Merdes et al., 1996). Furthermore, dynein–driven anti-parallel MT-MT sliding has also been demonstrated to be important for spindle fusion (Gatlin et al., 2009). These results identify dynein-driven MT-MT sliding as a necessary activity during spindle formation and maintenance.

How does dynein activity drive MT organization at the molecular level? Isolated dynein can generate forces to slide anti-parallel MTs in vitro and is sufficient to counteract kinesin-5 activity in vivo (Tanenbaum et al., 2013). Because this mechanism of dynein-driven sliding is restricted to anti-parallel MTs, how this observation is related to the focusing of parallel MT minus-ends at spindle poles is unclear (Goshima et al., 2005; Merdes et al., 2000; Morales-Mulia and Scholey, 2005). Furthermore, dynein’s role in spindle assembly likely requires its accessory factor dynactin (Echeverri et al., 1996; Merdes et al., 2000; Mitchison et al., 2005; Wittmann and Hyman, 1999)(though see (Raaijmakers et al., 2013)), suggesting that anti-parallel MT-MT sliding by isolated dynein may not be physiologically relevant within the spindle. Thus, the molecular mechanism for how dynein exerts forces within the spindle remains unknown.

Isolated dynein is not strongly processive, but its motility is greatly stimulated by its association with the ~ 1 MDa multi-subunit activating complex dynactin (McKenney et al., 2014; Schlager et al., 2014; Torisawa et al., 2014). This molecular interaction requires a third, coiled-coil adapter protein to mediate dynein-dynactin binding, forming a tripartite dynein-dynactin-activator complex (McKenney et al., 2014; Schlager et al., 2014). The most well-characterized coiled-coil activator, BicD2, strongly stimulates the formation of an ultra-processive dynein-dynactin-BicD2 co-complex (DDB) (McKenney et al., 2014; Schlager et al., 2014; Urnavicius et al., 2015). Upon activation, DDB moves robustly towards MT minus-ends, and was previously observed to strongly accumulate at MT ends in vitro (McKenney et al., 2014). Similarly, accumulations of dynein and dynactin at MT minus-ends generated by laser severing of spindle fibers have also been reported in vivo (Elting et al., 2014). During our studies of DDB motility in vitro, we observed that accumulations of motors at MT minus-ends were capable of generating robust forces along neighboring MTs. DDB accumulations were capable of simultaneously remaining bound to the minus-end of one MT, while generating pulling forces along another towards the minus-ends of a neighboring MT.

We have now studied this phenomenon in detail, and provide evidence that DDB accumulation at minus-ends is cooperative in nature, suggesting an interaction between motors, and is limited by the protofilament number of the MT lattice. Strikingly, minus-end accumulations of DDB can drive the robust sliding of MTs, without a preference for initial MT orientation. Multiple sliding events lead to the dramatic rearrangement of MTs and the focusing of multiple MT minus-ends together, similar to dynein’s functions in spindle assembly. At larger length-scales, DDB drives the formation and coalescence of MT asters in vitro, recapitulating an activity previously shown to be dynein-dependent in cell extracts (Gaglio et al., 1995; Verde et al., 1991). This activity is sufficient to cause the bulk contraction of millimeter scale MT networks, a phenomenon previously observed in *Xenopus laevis* extract (Foster et al., 2015). Thus, the cooperative accumulation of activated dynein motors at MT minus-ends provides a simple and robust mechanism to drive large-scale reorganizations of MTs (Belmonte et al., 2017). Our data provides a new molecular mechanism for how dynein fulfills its various roles during spindle assembly and in MT network reorganization in general.

## Results

### Processive Dynein-Dynactin Complexes Accumulate at MT Minus-Ends

To explore the kinetics of DDB accumulation at MT minus-ends, we created a ‘drop-in’ assay where the motor complex is added to an observation chamber during continuous acquisition, allowing the observation of the accumulation of DDB on microtubules over time (Fig. 1A, Movie S1). We used total internal reflection fluorescence (TIRF) microscopy to monitor DDB motility and accumulation along fluorescently-labeled MTs fixed to a coverglass. DDB molecules on the microtubule either moved processively along MTs toward the minus-ends or diffused on the MT lattice, as previously described (McKenney et al., 2014; Schlager et al., 2014) (Fig. 1A, B). Notably, individual processive molecules often did not dissociate when reaching the minus-end of the MT, but rather remained bound, leading to the accumulation of multiple motors at the minus-end (Fig. 1A, B, Movie S1). Quantification of the fluorescence signal in both the MT and DDB channels further confirmed that DDB accumulated at the ends of MTs (Fig. 1C), and the DDB intensity at individual MT ends reached a steady state during our observation period of ~ 5 min. (Fig. 1D).

**Figure 1:**
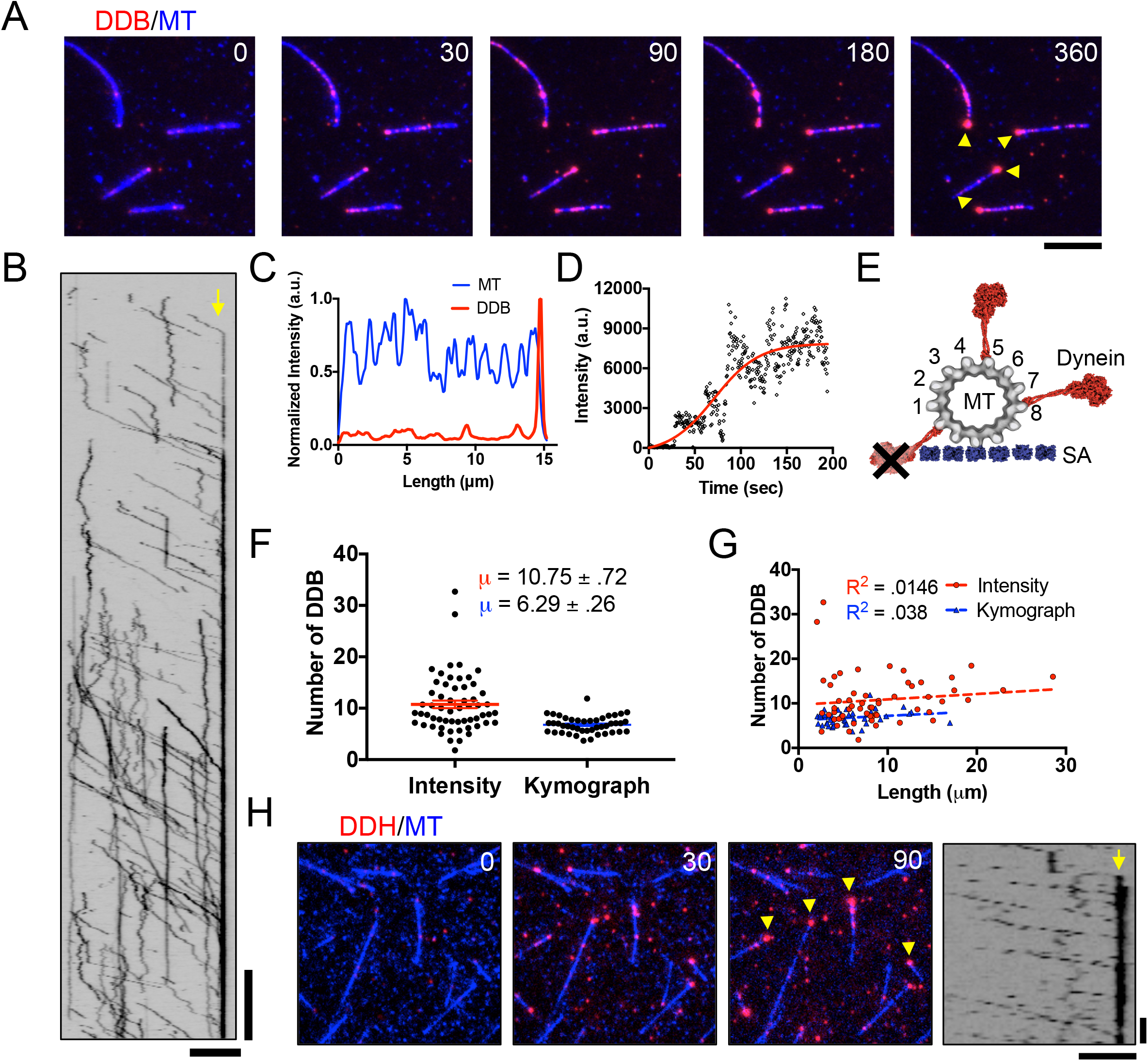
Accumulation of Processive Dynein-Dynactin Complexes at MT Minus-ends. **A)** TIRF-M images of DDB (red) accumulating at MT (blue) minus-ends. DDB accumulations are marked by yellow arrows. Scale bar: 2 μm, time is given in sec. **B)** Kymograph of DDB motility and accumulation on MT minus-ends. Minus-end denoted by yellow arrow. Scale bars: 2 μm and 30 sec. **C)** Quantification of MT (blue) and DDB (red) average intensities over time per pixel along a MT, normalized to the maximum intensity along the same line in the respective channel. **D)** Plot of minus-end DDB intensity over time. A 4-parameter logistic sigmoidal regression was fitted by least squares fit to the data (red), R^2^ = .795. **E)** Scale diagram of dynein motor domains (red) bound to a 13 pf MT. The MT is attached to a surface via streptavidin molecules (SA, blue), as in our assays. Not all protofilaments are available for dynein binding on the surface-bound 13-protofilament microtubule due to dynein’s large size. Note that the entire DDB complex is much larger than the dynein motor domain depicted. **F)** Quantification of the number of DDB complexes within minus-end accumulations using either fluorescent intensity analysis (red) or kinetic calculation methods (blue). N = 58 and 44 respectively, 2 independent experiments, see also Fig. S1. Data represented as mean ± SEM. **G)** Plot of number of DDB in minus-end accumulations vs MT length using either fluorescent intensity analysis at minus-ends (red), or using kinetic calculation methods (blue). N = 58 and 44 respectively, 2 independent experiments. **H)** DDH complexes (red) also form accumulations at MT (blue) minus-ends, denoted by yellow arrows. Right, Kymograph of processive DDH complexes accumulating at to MT minus-ends (yellow arrow). Scale bars: 5 μm and 10 sec.

Next, we aimed to quantify the number of DDB complexes within the steady-state accumulations at MT minus-ends. Due to the large size of the DDB complex (Chowdhury et al., 2015), and geometric constraints of surface-attached MTs, we estimated that ~ 7-8 MT protofilaments are accessible to the DDB complex at minus-ends in our assays (Fig. 1E). To quantify the numbers of DDB at minus-end accumulations, we used two methods to estimate the number of DDB molecules at steady state within the minus-end accumulations. First, we compared the integrated fluorescence intensity of the steady state minus-end accumulation to the intensity of single DDB molecules on the MT lattice, or nonspecifically bound to the glass near the MT (Fig. S1). Second, we used kymograph analysis to calculate the flux of DDB molecules into the minus-ends, as well as the off-rate of single DDB molecules from minus-ends (Fig. S1 and see below). Both methods yielded similar estimates of the number of individual DDB complexes within each minus-end accumulation, 10.75 ± 0.72 and 6.29 ± 0.25 complexes, respectively (Fig. 1F). This equates to approximately one to two DDB complex per MT protofilament, assuming most taxol-stabilized MTs in our preparation contain 12 protofilaments, and not all protofilaments are accessible to the motor due to the covalent linkage of the MT to the glass surface (Fig. 1E). Furthermore, the estimated number of DDB molecules in the minus-end accumulations did not correlate with MT length (Fig. 1F). These results argue that DDB molecules accumulate at MT minus-ends by prolonged association with single MT protofilaments.

To examine if minus-end accumulation was specific to dynein-dynactin complexes formed with the BicD2N adapter protein, we performed similar experiments with an orthogonal adapter molecule, Hook3, which has also been previously shown to mediate dynein’s interaction with dynactin, and activate processive dynein motility (McKenney et al., 2014; Olenick et al., 2016; Schroeder and Vale, 2016). Isolated dynein-dynactin-Hook3 complexes (DDH) displayed robust processive movement, as previously described, and noticeably accumulated at MT minus-ends similar to DDB complexes (Fig. 1H). Thus, we conclude that minus-end accumulation is an intrinsic property of activated, processive dynein-dynactin complexes, and does not depend on the type of adapter molecule that mediates dynein’s interaction with dynactin.

### Mechanism of Minus-End Accumulation

To explore the mechanism of minus-end accumulation, we first measured the rate of dissociation for individual DDB complexes at MT minus-ends (Fig. 2A, D, E, movie S2). We found that the resulting dwell time distribution was best fit with a biphasic decay, suggesting two types of dissociation events at minus-ends (Fig. 2D, E). The fit revealed a shorter dwell time of 9.39 s and a much longer dwell time of 54.8 s, with 56.76% of the molecules displaying the faster dissociation time (Fig. 2E). Thus, single molecules of DDB exhibit both fast and slow dissociation from MT minus-ends, with a preference for the former in our assay conditions.

**Figure 2:**
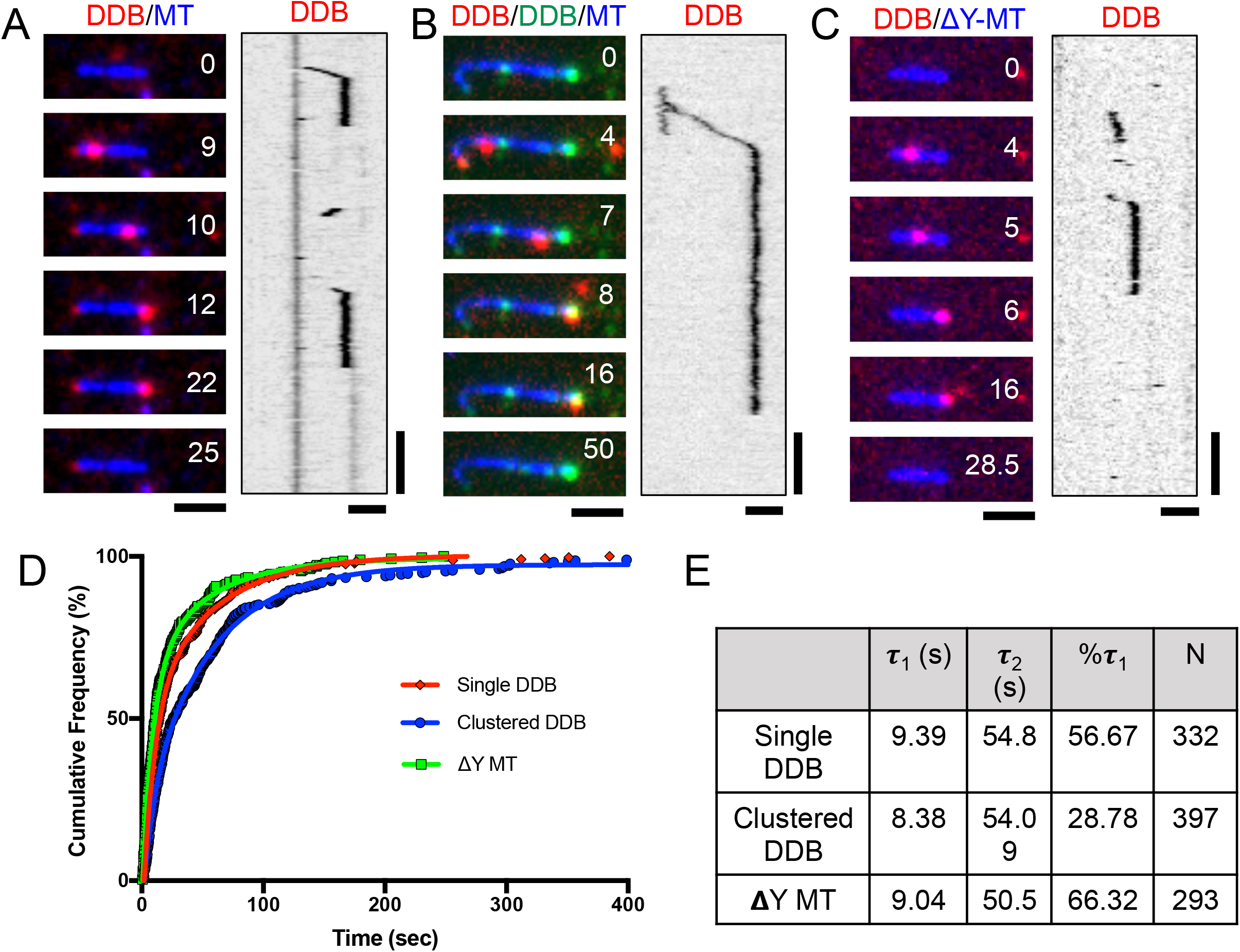
Single Molecule Analysis of DDB Dwell Times at MT Minus-Ends. **A)** Representative TIRF-M images and associated kymographs of DDB (red) behavior at MT (blue) minus-ends. **B)** Representative images and associated kymograph of single molecule spiking experiments at ~1 nM SNAP-TMR DDB (red) and ~30 nM SNAP-488 DDB (green) dwelling at MT minus-ends (blue). **C)** Representative images and associated kymograph of DDB (red) dwelling at the ends of carboxypeptidase treated MTs (blue). Time is given in sec. for A-C. Scale bars: 2 μm and 15 sec. **D)** Cumulative frequency plots of DDB dwell times from previous conditions. Solid lines are two-phase exponential regressions to the data. **E)** Table summarizing the parameters of DDX cumulative frequency graphs, including the characteristic dwell time (τ) of short-dwelling (τ_1_), long-dwelling (τ_2_) populations, percentage of molecules in the short-dwelling population (%τ_1_), number of molecules measured (N), and goodness of fit (R^2^). All data from 2-3 independent experiments.

To gain insight into how minus-end dissociation rates might be affected by other motors within a minus-end accumulation, we next performed single molecule spiking experiments with differentially labeled DDB. We mixed together a ratio of SNAP-488 and SNAP-TMR labeled DDB complexes such that all MTs in the chamber contained accumulations of SNAP-488 DDB at their minus-ends, while only individual molecules of SNAP-TMR DDB were observed to bind and move along MTs (Fig. 2B, Movie S2). We then measured the dwell times of single SNAP-TMR DDBs within the context of the SNAP-488 minus-end accumulations. Strikingly, individual SNAP-TMR DDBs appeared to dwell much longer within minus-end accumulations of SNAP-488 DDB (Fig. 2B, D, E). The same effect was observed when SNAP-647 DDB complexes were used to form minus-end accumulations, arguing against any effect of the SNAP dye used to label the DDB complexes. Quantitative analysis revealed that individual dwell times were not changed significantly, but there was nearly a two-fold shift in the population towards longer dwell times (Fig. 2B, D, E). These results show that individual DDB molecules shift their behavior at MT minus-ends depending on the presence of other DDB molecules at the MT end. Such a change in behavior may suggest a cooperative interaction between DDB molecules at MT minus-ends that shifts equilibrium between two DDB dissociation rates towards longer dwell times.

DDB contains two distinct types of MT-binding domains (MTBD). Dynein’s MTBD is located within the motor domain, and binds directly to the tubulin lattice (Carter et al., 2008; Redwine et al., 2012). The second MTBD, located at the N-terminus of the p150^*Glued*^ subunit of dynactin, binds to the disordered tubulin tail domains (Culver-Hanlon et al., 2006; Lazarus et al., 2013; McKenney et al., 2016; Peris et al., 2006; Wang et al., 2014). In principle, the association of DDB to minus-ends could be mediated by one or both MTBDs. To distinguish between these possibilities, we treated taxol-stabilized MTs with carboxypeptidase A (CPA) to remove the C-terminal tyrosine residue of α-tubulin (Webster et al., 1987), which has been previously shown to greatly diminish the interaction of the p150^Glued^ MTBD with the MT lattice, but has comparatively little effect on the interaction of dynein with MTs (McKenney et al., 2016). This treatment did not significantly change the DDB dwell time distribution as compared to WT MTs, suggesting that engagement of the p150^Glued^ MTBD with the MT lattice is not required for extended complex dwelling at MT minus-ends (Fig. 2C, D, E, Movie S2). We conclude that DDB complexes accumulate cooperatively on MT minus-ends through interactions of the dynein MTBD with the MT lattice.

### MT Reorganization by Minus-End Accumulations of DDB

The ability to produce force along one MT while remaining bound to another is the basis for MT reorganization driven by molecular motors in cells. Isolated non-processive dimeric dynein can produce force from within MT-MT overlaps, possibly due to the low coordination and high flexibility between its two motor domains (Amos, 1989; Tanenbaum et al., 2013). However, we noted that processive DDB did not obviously produce force between overlapping MTs in our assays (Fig. 3A), reflecting a distinction between isolated dynein and dynein in complex with dynactin. Instead, forces between adjacent MTs were clearly visible once the minus-end DDB accumulation on one MT contacted an adjacent MT (Fig. 3A).

**Figure 3:**
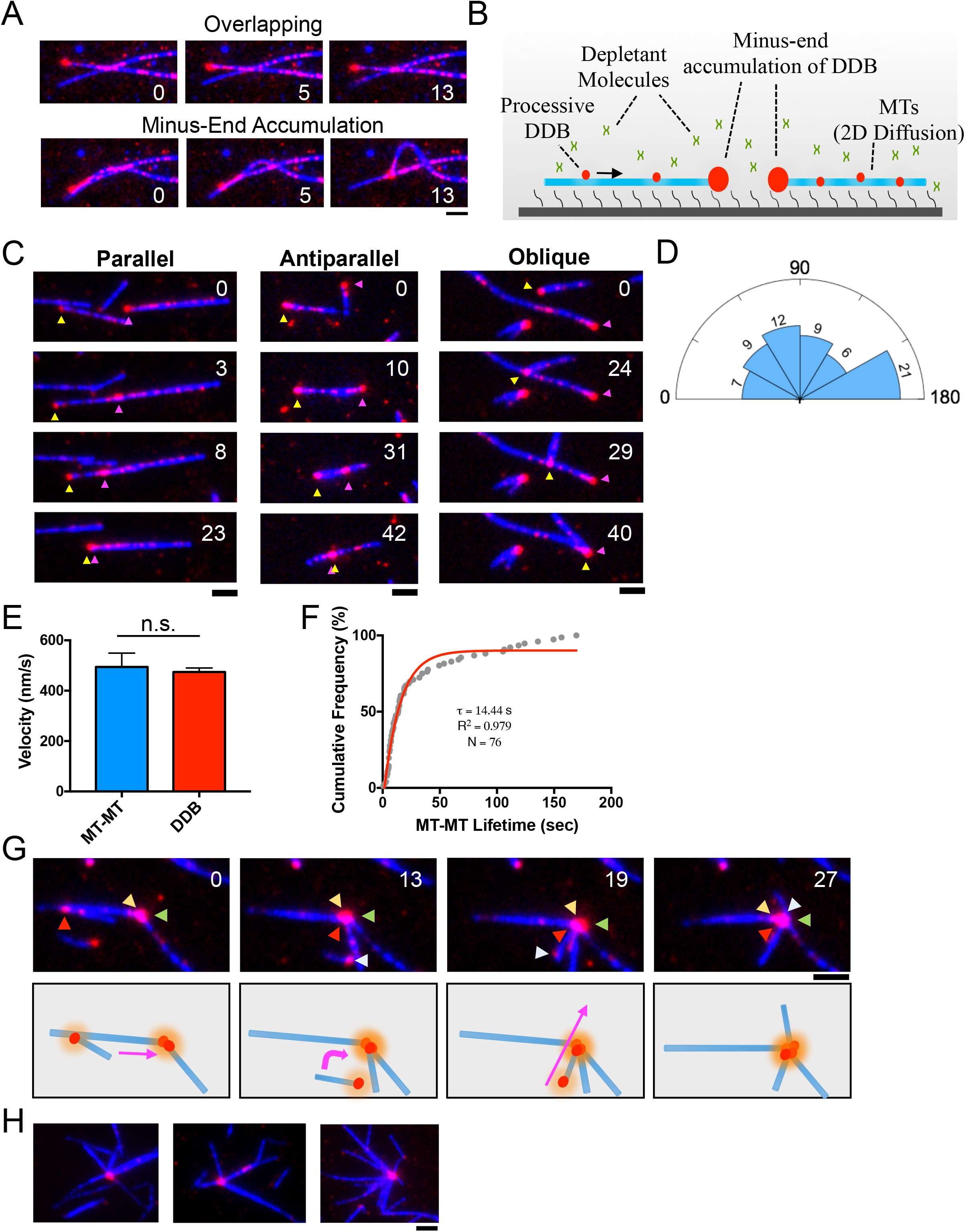
Minus-End Clustering Driven by Minus-End Accumulations of DDB. **A)** TIRF-M images of two separate events on the same set of loosely attached, overlapping microtubules (blue) without (top), and with (bottom), contact with the minus-end DDB cluster. Time is in sec., scale bar 2 μm. **B)** Schematic of TIRF experiments involving microtubules held in the TIRF field non-covalently using depletion forces. In this system, MTs are free to move in two dimensions along the coverslip surface. **C)** Representative TIRF-M images of parallel, antiparallel and oblique sliding by DDB minus-end clusters. MT minus-ends are inferred from DDB accumulation location, and designated by magenta and yellow arrows. **D)** Rose diagram representing the relative distribution of initial angle of MT-MT sliding. Endpoints (0, 180°) were integrated into their nearest bins. The number of sliding events in each bin are indicated. Bin size = 30 degrees. N = 64. **E)** Bar graph displaying the mean velocity of MT-MT sliding (blue) (474.1 ± 16.33 nm/s, N = 107, 2 independent experiments) in comparison to velocity of individual DDB molecules along MTs (red) (494.1 ± 55.09 nm/s, N = 43, 3 independent experiments). There is no statistical difference between the two conditions of DDB movement (P = 0.644, two-tailed T-test). Data represented as mean ± SEM. **F)** Cumulative frequency plot of MT-MT foci life times. Solid red line is the exponential regression fit to the data. Data composed of events where only two MT minus-ends are brought into contact by DDB motility and subsequently dissociate. τ = 14.44 s, N = 76, R^2^ = .979, 3 independent experiments. **G)** Top: TIRF-M images showing multiple MT-MT sliding events forming a mini-aster. Arrows label minus-ends of unique microtubules, inferred from DDB accumulation. Time is given in sec. Bottom: Schematic of MT movements observed in the movie. Colored arrows indicate direction of MT movement. **H)** More examples of mini-asters observed during the reorganization assay. Scale bars: 2 μm.

In the previous type of assay, MTs are bound tightly to the cover glass through near covalent biotin-streptavidin linkages between the MT lattice and the glass surface, greatly constraining the DDB-driven movement of MTs relative to one another. To fully explore the ability of DDB to drive MT-MT movements, we modified this assay to allow complete freedom of movement while keeping the MTs near the coverglass and within the TIRF illumination depth. We took advantage of the depletion force exerted by the addition of inert polymer methylcellulose (Henkin et al., 2014), which prevents MTs from diffusing away from the coverglass, but still allowed freedom of movement in two dimensions along the glass surface (Fig. 3B, Movie S3). This modification of the standard single molecule assay revealed the full capabilities of DDB-driven MT-MT sliding.

Using this new assay, we observed that minus-end accumulations of DDB could robustly drive the sliding of MTs in nearly any orientation, including parallel, anti-parallel, and oblique angles (Fig. 3C, D, Movie S3). Quantification revealed no obvious constraints on the angle of sliding driven by DDB (Fig. 3D). MT-MT sliding velocity was identical to the speed of individual DDB complexes moving along the MT lattice (Fig. 3E). Sliding always ended when the two minus-ends of the respective MTs (marked by high DDB signal) came within a diffraction-limited distance from one another, and quantification of the lifetime of the interaction between two minus-ends revealed a prolonged interaction lasting ~ 14 seconds (Fig. 3F). Multiple types of these sliding events often resulted in the accumulation of several individual MT minus-ends held together at the center of small asters (Fig. 3G, H, Movie S4), which tended to accumulate over time during the assay. Similar sliding events and small aster formation was observed when the experiments were repeated with DDH complexes, indicating that this phenomenon is not unique to the BicD2N adapter protein, but rather a molecular property of activated, processive dynein-dynactin complexes (Fig. S2B,C,D). Together, this data demonstrates that minus-end accumulations of DDB drive the robust sliding of MTs in any orientation resulting in two or more minus-ends brought into close proximity and held together for significant periods of time, and that activated, processive dynein-dynactin complexes are sufficient to induce the formation of small MT asters.

### Self-organization of MTs is Driven by Minus-end Accumulations of DDB

Because the previous analysis was performed on taxol-stabilized MTs, we wondered how DDB complexes interact with the minus-ends of dynamic MTs, as found in cells. We reconstituted dynamic MTs in vitro and observed DDB complexes moved along the full length of the lattice, and strongly accumulated at the dynamic minus-ends (Fig. 4A, Movie S5). Further, accumulations of DDB on dynamic minus-ends were capable of driving MT-MT interactions (Fig. 4B, Movie S5), similar to what we observed with taxol-stabilized MTs (Fig. 3). These interactions often resulted in the focusing of two or more dynamic minus-ends together (Fig. 4B), a function that we note is one of dynein’s key roles during mitotic spindle assembly (Goshima et al., 2005; Heald et al., 1996; Morales-Mulia and Scholey, 2005).

**Figure 4:**
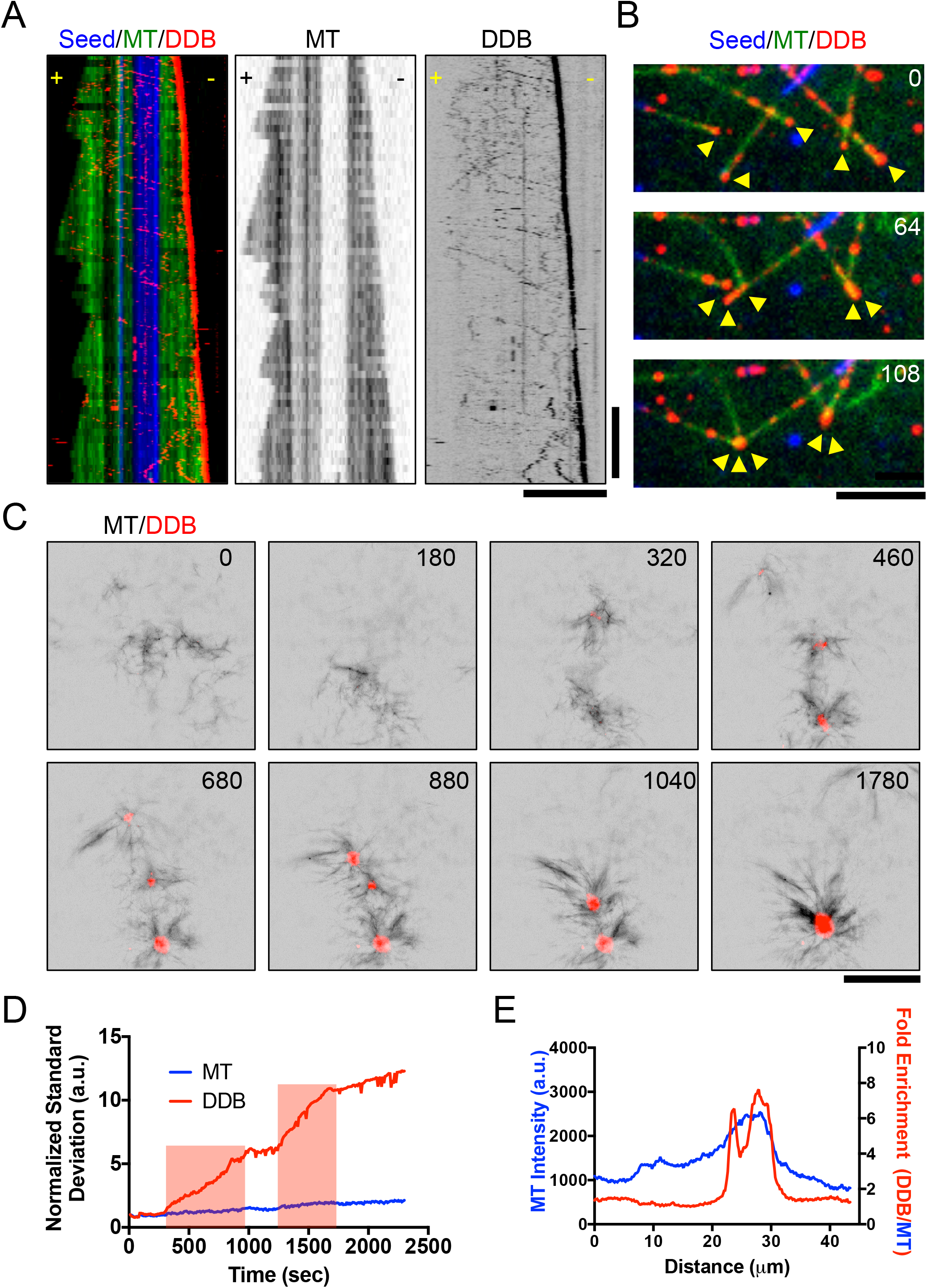
Minus-End Accumulations of DDB Drive Mesoscale MT Reorganization. **A)** Three-color kymograph of processive DDB (red) reveals strong accumulation at the growing minus-ends of dynamic MTs (green). Dynamic MT ends grown from GMPCPP-stabilized seeds (blue). Right: Individual fluorescent channels are reproduced for dynamic MTs and DDB for clarity. Scale Bars: 5 μm and 2 minutes. **B)** Example of DDB accumulations (red) on growing MT minus-ends (green) driving MT-MT sliding to form minus-end clusters (arrows). Note that MT-MT sliding does not occur at MT crossover points not in contact with minus-end DDB clusters. Time in seconds. Scale Bar: 5 μm **C)** Images of DDB-driven reorganization of growing microtubules (black) into asters with DDB (red) centers. Multiple asters coalesce into a single larger structure. Scale bar: 10 μm. Time in seconds. **D)** Plot of intensity contrast of both MT (blue) and DDB (red) channels expressed as standard deviation over time. Red bars indicate periods of merging between two asters. **E)** Line scan plot of MT intensity (blue) across an aster and the fold increase of DDB fluorescence intensity divided by MT fluorescence intensity.

In cells, the collective action of MT motors drives the self-organization of the MT cytoskeleton. Our previous assays demonstrated that minus-end accumulations of DDB complexes are sufficient to sort and organize MTs by sliding until minus-ends are brought together. To understand how DDB accumulations at minus-ends could drive MT-MT interactions at a mesoscale level, on the order of hundreds of MTs, we studied the behavior of DDB in solutions of growing MTs. When 4 μM DDB was incubated in solutions of growing MTs (40 μM tubulin, 2 μM taxol, 2mM GTP), we observed the formation and coalescence of micron-size asters over time (Fig. 4C, Movie S6). DDB signal accumulated strongly at the center of the MT asters during their formation (Fig. 4C), indicating the asters formed by DDB-driven collection and focusing of MT minus-ends. Nearby asters often coalesced and fused into a single larger aster (Fig. 4C), demonstrating a net contractile force produced by DDB in this system. Analysis of the image contrast, a measure of organization (Hentrich and Surrey, 2010), in the DDB channel showed the amount of contrast greatly increased at times corresponding to aster merging, and plateaued after a fusion event indicating DDB accumulation reaches steady state after the fusion event (Fig. 4D). Line-scan analysis revealed that DDB intensity at the center of the asters was enriched relative to the MT signal (Fig. 4E), indicating that the DDB foci accumulate specifically in the center of the asters, presumably at MT minus-ends. Cumulatively, these data demonstrate that processive DDB complexes cooperatively accumulate at growing MT minus-ends where they drive MT self-organization, focusing of MT minus-ends into asters, and the coalescence and fusion of asters.

### DDB Activity Drives Macroscale Contractile Stresses in MT Networks

The previous data demonstrated that DDB motor activity localized at MT minus-ends drives MT reorganization by clustering minus-ends together into MT asters. The coalescence and fusion of these asters reveals a net contractile stress exerted by DDB motor activity at MT minus-ends. Previous work in *Xenopus* egg extract revealed a dynein-dependent stress that drives the bulk contraction of millimeter-scale stabilized MT networks (Foster et al., 2015). These dynein-driven contractile stresses were hypothesized to be related to dynein’s role in mitotic spindle pole focusing, spindle healing, and in spindle fusion. The bulk contraction observed in extracts was driven by MT aster formation and coalescence, similar to what we observe in purified solutions of DDB and MTs (Fig. 4). We therefore set out to determine if purified DDB could recapitulate bulk MT network contraction, independent of other cellular factors found in *Xenopus* egg cytoplasm.

We combined various concentrations of purified, TMR-labeled DDB complexes with taxol stabilized, Atto- or Alexa-647-labeled MTs in microfluidic devices measuring 125 μm (H) × 0.9 mm (W) × 18 mm (L). In this system, the initially diffuse MT network exhibited spontaneous bulk contraction on the millimeter length scale, strongly resembling the bulk contraction previously observed in *Xenopus* extract (Foster et al., 2015) (Fig. 5A, Movie S7). Imaging of the contracting network at higher magnification revealed the network was organized into aster-like structures with DDB fluorescence concentrated in the aster centers (Fig. 5B, Movie S8), consistent with the network structures observed previously in *Xenopus* extracts (Foster et al., 2015). We quantified the contraction dynamics by measuring the width of the network as a function of time, *W*(*t*). These measurements were then converted into the fraction the network has contracted as a function of time, *ϵ*(*t*), using the relation,

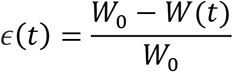

where *W_0_* is the initial width of the network. These *ϵ*(*t*) curves were well described by a saturating exponential function,

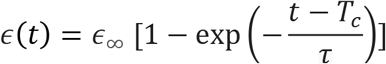

where *τ* is the contraction timescale, *ϵ*_∞_ is the final fraction contracted, and *T_c_* is a lag time. Fitting the measured *ϵ*(*t*) curves allowed these parameters to be measured for each experiment. When the concentration of DDB was increased, the contraction timescale was found to decrease, with only a slight increase in the final fraction the network contracts (Fig. 5C). This result is reminiscent of previous dynein inhibition experiments in extract, and in qualitative agreement with the previously proposed active fluid model (Foster et al., 2015)(Fig. 5C, D). The model predicts that the strength of contractile stresses, and thus the contraction timescale of the active fluid should depend on the concentration of motors, while the final density of the MT network should remain unchanged. Thus, a mixture of purified processive dynein-dynactin complexes and stabilized MTs is sufficient to recapitulate the bulk MT network contraction observed in *Xenopus* egg cytoplasm.

**Figure 5:**
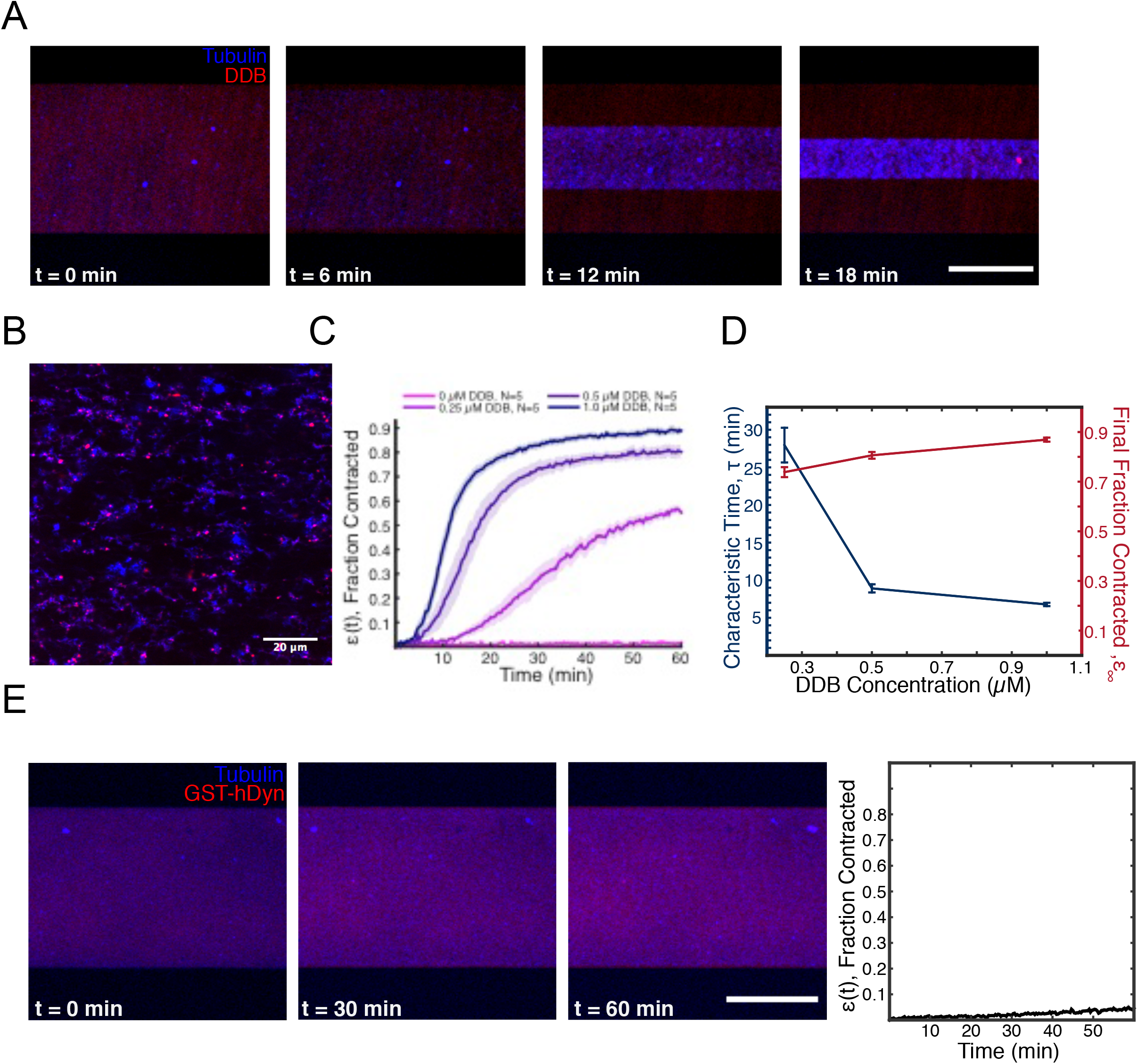
DDB Drives Bulk Contraction of Macroscale Networks of Microtubules. **A)** Representative images of a microfluidic containing a network of taxol-stabilized microtubules (blue). The MT network contracts over time due to forces exerted by DDB (red). Scale bar: 500 μm. **B)** Higher magnification image of the contracting network of microtubules (blue), showing aster-like structures, with DDB (magenta) localizing towards the interior. Scale bar: 20 μm **C)** Average fractional network contraction as a function of time for varying DDB concentrations (mean ± s.e.m). **D)** Characteristic time, τ, and final fraction contracted, ε_∞_, from fits to ε(t) curves as a function of DDB concentration. **E)** Representative images of a microfluidic channel containing a network of taxol-stabilized microtubules (blue) and purified GST-hDyn (red). Note the network shows little contraction over much longer timescales than in (A). Right: Quantification of average fractional network contraction as a function of time.

A recombinant, minimal dimeric human dynein construct, GST-hDyn, was previously shown to be capable of bundling and sliding anti-parallel MTs in vitro, and antagonizing Eg5 activity within the mitotic spindle in vivo (Tanenbaum et al., 2013). Because this construct is not strongly processive (McKenney et al., 2014; Torisawa et al., 2014; Trokter et al., 2012), it cannot accumulate at MT minus-ends as we observe for DDB in vitro. Rather than generating forces at MT ends, GST-hDyn is thought to generate forces from within a MT-MT overlap using its two uncoordinated motor domains (DeWitt et al., 2012; Qiu et al., 2012; Tanenbaum et al., 2013). As a result, the sliding activity of GST-hDyn is restricted to anti-parallel MTs, whereas minus-end accumulations of DDB are not restricted by MT geometry (Fig. 3C). To test if processivity is important for dynein-driven MT network contraction, we incubated GST-hDyn with mixtures of MTs and observed minimal network contraction, even after much longer time scales than we investigated for DDB (Fig. 5E, Movie S7). We conclude that cooperative accumulation of activated dynein-dynactin complexes at MT minus-ends provides a simple and robust mechanism for contractile stress generation within MT networks, and suggest this mechanism can account for dynein-dependent contractile stress generation within the mitotic spindle.

## Discussion

### Behavior of Individual Dynein-Dynactin Complexes at MT Minus-ends

Microtubule motors drive the self-organization of MT-based structures within cells. How single types of motors, as well as ensembles of different types of motors cooperate to produce such organization is an area of intense interest. Previous work has provided molecular insight into how several classes of kinesin motors function in MT organization, particularly during mitotic spindle self-assembly (Cross and McAinsh, 2014). In addition to various kinesins, cytoplasmic dynein plays several roles in intracellular MT organization, and is critical for normal bipolar spindle assembly (Heald et al., 1996; Merdes et al., 1996; Raaijmakers et al., 2013; van Heesbeen et al., 2014). Dynein activity is necessary to collect, focus, and anchor MT minus-ends into the spindle pole (Goshima et al., 2005; Maiato et al., 2004; Morales-Mulia and Scholey, 2005; Raaijmakers et al., 2013). In addition, dynein is required for bipolar spindle morphology, in part through antagonizing kinesin-5 and kinesin-12, which act on antiparallel MTs (Cross and McAinsh, 2014; Drechsler and McAinsh, 2016; Drechsler et al., 2014; Kapitein et al., 2005; Sharp et al., 1999; Sturgill and Ohi, 2013; van den Wildenberg et al., 2008). The molecular mechanism for how dynein fulfills these roles in spindle assembly remains unknown, and our results provide new insight into how dynein motor activity can be harnessed for MT organization.

Here, we have analyzed the single molecule and bulk behavior of activated, processive dynein-dynactin complexes at length-scales that span several orders of magnitude. At the nanoscale, we find that stable dynein-dynactin complexes, formed using two distinct adapter molecules, accumulate cooperatively at the minus-ends of single MTs, the natural endpoint of any dynein-based movement in cells. The cooperativity we observe in DDB accumulation may reflect molecular interactions between individual DDB complexes at MT ends. The molecular basis of such an interaction remains to be determined, but we speculate that it could arise through motor domain interactions between adjacent DDB complexes in trans, akin to previously observed interactions between motor domains within a single dynein homodimer (Fig. 6) (Amos, 1989; Kon et al., 2011; Torisawa et al., 2014; Toropova et al., 2017). It is also possible that DDB’s that reach the MT minus-end could impart structural changes to the MT lattice that favor further DDB retention. Indeed, dynein motor activity has been shown to affect MT plus-end dynamics in vitro (Laan et al., 2012) (Hendricks et al., 2012) and further work will be needed to determine if DDB affects minus-end dynamics.

**Figure 6:**
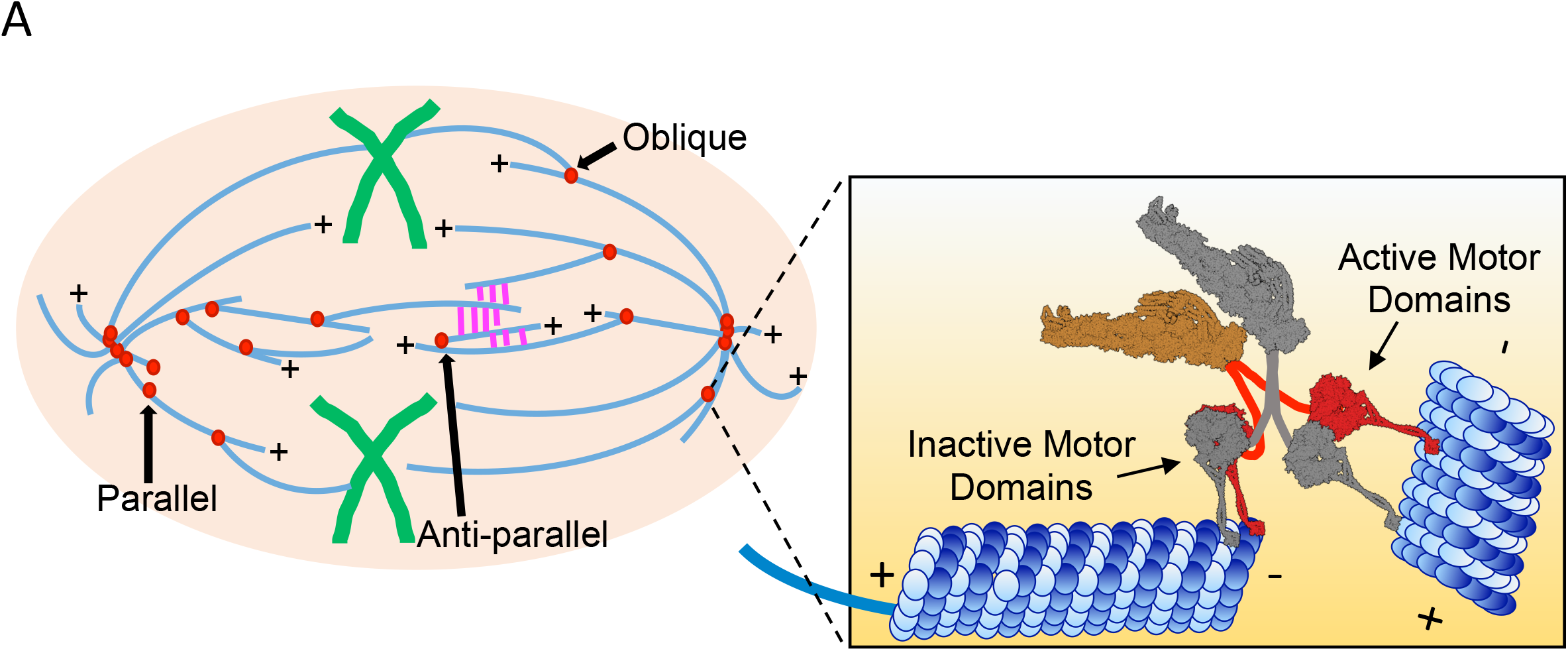
Model For Cooperative Accumulation of DDB Complexes At MT Minus-ends That Drive MT Organization Within the Bipolar Mitotic Spindle. Different orientations of MT-MT sliding are required at topologically unique locations within the mitotic spindle. Parallel sliding by minus-end DDB complexes (red) is important for spindle pole focusing while anti-parallel sliding resists kinesin forces (magenta) within the spindle midzone. Oblique interactions are likely important in integrating MTs into the spindle network (Elting et al., 2014; Redemann et al., 2017; Sikirzhytski et al., 2014). Inset: DDB complexes accumulate on MT minus-ends. We propose that individual complexes adopt inactive confirmation, possibly through trans interactions with neighboring DDBs (gray), providing an anchor to the MT minus-end. Single motor domains from each DDB complex are free to bind, and produce force towards the minus-ends of adjacent MTs along driving minus-end directed MT-MT sliding which produce net contractile forces within MT networks.

The number of DDB complexes at MT ends reaches steady state, suggesting an equilibrium between on and off rates at the minus-ends. Our measurements provide evidence that the number of DDB complexes within minus-end accumulations is limited by the binding site for the dynein motor domain on each MT protofilament. We note the size of the accumulations do not change appreciably during periods of increased DDB flux into the minus-end (Fig. 1B), further suggesting limited binding sites at MT ends. Because DDB accumulation does not require an interaction between p150^Glued^ and the MT surface, we favor a model whereby individual DDB complexes remain bound to minus-ends via one or both of their dynein motor domains.

### Minus-End DDB Reorganizes MT Networks

Motor forces exerted from within overlaps drive most MT-MT sliding studied to date. Because of this, the geometry of sliding is typically limited by the intrinsic ability of the motor to move in one direction along the lattice. Here, we find that DDB produces force on adjacent MTs predominately from accumulations at MT minus-ends, allowing a much greater degree of freedom to drive MT-MT sliding regardless of MT orientation. The result of all DDB catalyzed sliding events, without a bias for initial MT orientation, is the coalescence of MT minus-ends together. This feature makes MT-MT sliding generated by DDB unique among MT motors reported, and may be a consequence of the relatively uncoupled mechanochemistry and flexibility of dynein’s two motor domains (DeWitt et al., 2012; Qiu et al., 2012).

The ability to slide parallel as well as anti-parallel MTs may provide direct insight into dynein’s known roles within different parts of the mitotic spindle (Fig. 6). In the spindle mid-zone, enriched in anti-parallel MTs, dynein activity is thought to antagonize kinesin-5 and kinesin-12 dependent extensile sliding (Florian and Mayer, 2012; Mitchison et al., 2005; Tanenbaum et al., 2008; Tanenbaum et al., 2009; van Heesbeen et al., 2014). In contrast, at the spindle poles, where parallel MTs predominate, dynein activity is important to collect and focus MT minus-ends and anchor them to the spindle pole (Goshima et al., 2005; Maiato et al., 2004; Morales-Mulia and Scholey, 2005; Raaijmakers et al., 2013). Importantly, the mechanism of sliding described here is distinct from that previously reported for dimerized minimal dynein constructs, and likely explains why these constructs were unable to rescue focusing of parallel MTs at spindle poles (Tanenbaum et al., 2013). In addition, accumulations of dynein-dynactin at nascent MT minus-ends, produced by laser severing, have been demonstrated to capture neighboring MTs at oblique angles and drive poleward parallel sliding to repair spindle architecture (Elting et al., 2014; Sikirzhytski et al., 2014). The mechanism of dynein-dependent MT-MT sliding we present here provides a simple and robust model for how dynein activity can fulfill these diverse functions within the mitotic spindle.

How do accumulations of dynein-dynactin produce force along adjacent MTs, while maintaining attachment to the minus-end? The cooperative mechanism of accumulation may provide a clue, as it suggests molecular interactions among motors within the accumulation. We hypothesize that each motor may remain bound to the minus-end via a single motor domain, while its partner motor domain remains free to bind and produce force along adjacent MTs. Because the motor domains of dynein are not strongly coupled, and individual dynein dimers composed of a single active motor domain can still move processively in vitro (Cleary et al., 2014), we hypothesize that the force generated from minus-ends may be the sum of many individual motor domains working independently from each other. We speculate that the minus-end bound motor domain may be held in a mechanically inactive, strongly MT bound state, through trans interactions with neighboring motors. Such a conformation could provide a stable tether to the minus-end of one MT, while allowing the free motor domain to produce force along an adjacent MT. However, our observation that DDBs remain accumulated at the growing minus-ends of dynamic MTs suggests that the motors within the accumulation are competent to rapidly switch between inactive, and actively processive states.

Because we observe two populations of off-rate for single DDBs, we suggest that the DDB complex samples at least two conformational states upon reaching the minus-end, and accumulation of more complexes favors one state over the other. The molecular basis for these two states remains to be determined, but we note that even the shorter dwell time that we observe at minus-ends is at least 20-fold longer than dwell times calculated in computer simulations to be effective in focusing of k-fiber minus-ends during spindle assembly (Goshima et al., 2005). This study concluded that the dwell time at minus-ends plays a critical role in determining if a molecular motor will perform well in focusing MT ends, an observation that our experimental data supports.

### Minus-End Accumulations Provide Contractile Forces for MT Self-Organization

We have also examined the emergent behavior of systems composed of MTs and DDB. At a mesoscale level, mixtures of growing MTs and DDB form micron-sized asters that fuse together via contractile forces exerted across tens of microns. Aster fusion is presumably driven by distal MT ends contacting the DDB core of another aster, and is reminiscent of dynein-driven aster and spindle fusion in *Xenopus* extracts (Gatlin et al., 2009). Previous in vitro studies have demonstrated similar aster formation using artificially multimerized kinesin motors (Nedelec et al., 1997; Surrey et al., 2001), or kinesin motors that naturally contain a second MT-binding site (Hentrich and Surrey, 2010). Aster formation in cell extracts has been shown to be dynein-dependent (Gaglio et al., 1995; Verde et al., 1991). Here we have shown that the native, dimeric DDB complex drives aster formation by generating force exclusively from small accumulations at MT ends, an important distinction from previous results with various kinesin motors. Further, DDB can slide and reorganize MTs in any orientation and possesses motor velocity and processivity large enough overcome the speed of minus-end growth. The high velocity and extreme processivity of DDB may make the motor complex particularly well-suited to reach MT ends compared with much slower velocity and limited processivity of kinesin-5 (Kapitein et al., 2008; Kwok et al., 2006) or kinesin-14 (Fink et al., 2009; Hentrich and Surrey, 2010).

Remarkably, we find that purified DDB complex is sufficient to completely recapitulate the bulk contraction of MT networks, at millimeter length scales, as observed previously in *Xenopus* extract (Foster et al., 2015). The contractile stresses exerted by dynein in mitotic extracts are likely to be related to the mechanisms used to cluster MT minus-ends during aster formation and spindle pole focusing in vivo. In cells, the mitotic molecules kinesin-14 (Goshima et al., 2005), NuMA (Merdes et al., 1996), and Lis1 (Moon et al., 2014; Raaijmakers et al., 2013; Tai et al., 2002) have been shown to be essential for the formation of asters and spindle poles. However, our results reveal that in vitro, dynein motor activity in isolation recapitulates these activities, raising new questions about the role of these proteins in cells. Kinesin-14 and dynein perform semi-redundant functions during spindle assembly (Goshima et al., 2005), and our work suggests that unbiased angle of sliding could be a key difference that differentiates dynein from kinesin-14 function.

In our assays, we used the well-characterized BicD2N adapter molecule to mediate the interaction between dynein and dynactin, but BicD2N has no known role in mitosis in vivo. We hypothesize that NuMA could fill a similar role of mediating the dynein-dynactin interaction, leading to dynein motor activation and accumulation at MT minus-ends. NuMA’s own MT-binding domain (Haren and Merdes, 2002) could serve to enhance minus-end clustering by further augmenting motor dwell time at minus-ends or by providing another tether to the MT lattice during MT-MT sliding. Finally, the dynein regulatory factor LIS1, which is required for normal spindle assembly and locks dynein in a strongly MT-bound state (Huang et al., 2012; McKenney et al., 2010), also accumulates with DDB at minus-ends in vitro (Gutierrez and McKenney, 2017). LIS1’s role at minus-ends remains to be examined, but it may further strengthen the DDB affinity for MT ends, leading to longer lasting interactions between MTs. Further experiments reconstituting the mitotic environment in vitro will provide insight into the role of these molecules and processes.

In summary, our work provides a mechanism for dynein-driven contractile forces in a MT network and proposes a model for how dynein sorts and organizes MTs at spindle poles. We propose that the contractile stresses involved in aster formation, poleward movement of k-fibers, and spindle pole focusing, are generated by clusters of activated, processive dynein at MT minus-ends.

## Author Contributions

RJM and DJN conceived the project. RT and PJF performed the experiments and analyzed the data. All authors wrote the manuscript and assembled the figures.

## Acknowledgements

Thanks to Elizabeth Christine Paz for generating the original macros for image analysis. We thank members of the Ori-McKenney lab for critical input during the project and Christina Hueschen and Sophie Dumont for valuable advice and feedback. We thank Bryce Ackermann for help with molecular structures. This work was supported by the Kavli Institute for Bionano Science and Technology at Harvard University, National Science Foundation grant DMR-0820484 to DJN, and startup funding provided by UCD to RJM.

---

**Figure S1:**
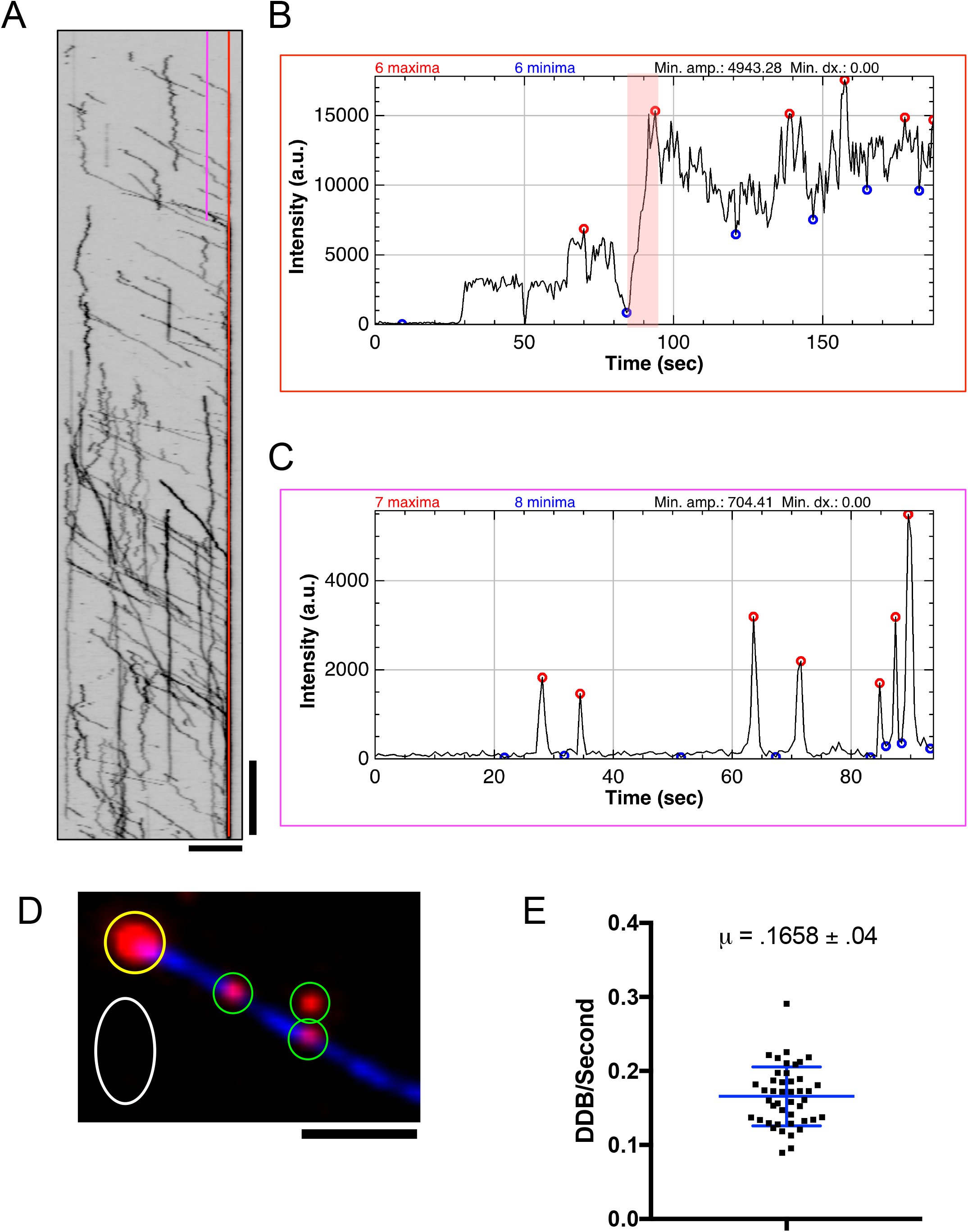
Determination of Numbers of DDB Molecules at MT Minus-Ends. **A)** A kymograph of a microtubule accumulating DDB at its minus-end (reproduced from Fig. 1B). A line scan was drawn along the region corresponding to the MT minus-end (red). The earliest peak of the fluorescence intensity plateau along this line scan was used as a fiducial for when the minus-end accumulation reaches saturation. A second line scan (pink) extending as far as the DDB intensity plateau fiducial in the Y-axis was drawn further into the MT as close to the minus-end as possible without interference from the growing minus-end accumulation. **B)** Intensity plot of the microtubule minus-end (red line in kymograph (A)). The earliest peak of the fluorescence intensity plateau as indicated by the red highlight was used as a fiducial for when a minus-end reaches saturation. **C)** Intensity plot in the microtubule lattice used to count the number of DDB that enter the minus-end accumulation until saturation. Every peak intensity on this new line scan corresponds to a single DDB crossing the point in space. Red circles indicate peaks and correspond to individual DDB. **D)** Example of intensity analysis for numbers of DDB at MT ends. Yellow, white, and green circles indicate ROIs for minus-end accumulations, background fluorescence normalization, and single molecules, respectively. Right: Unannotated image for clarity. Scale bar: 2 μm. **E)** Plot of the rate at which DDB enter the minus-end. μ = .1658 ± .04. N = 44.

**Figure S2:**
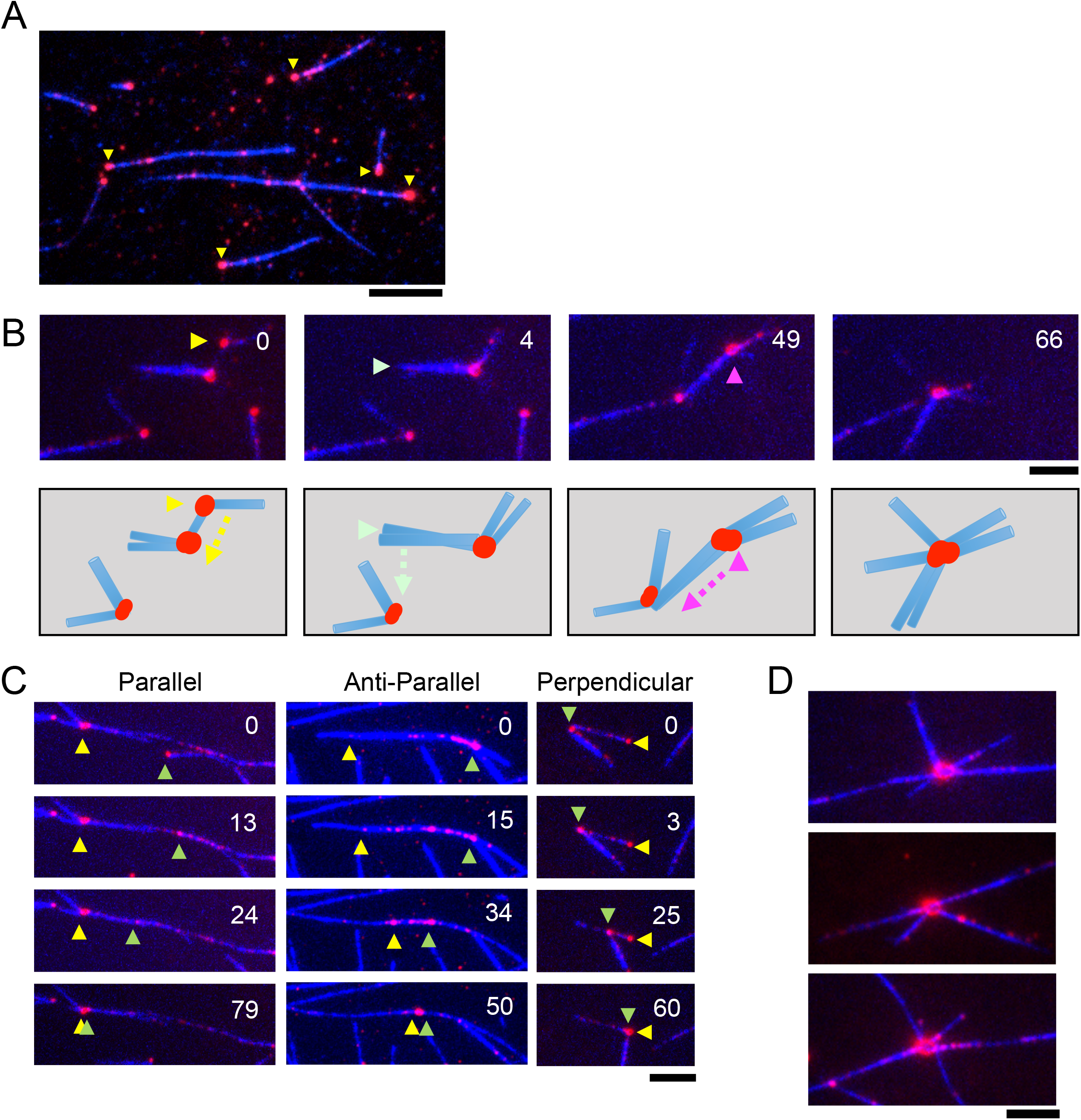
Minus-End Accumulations of DDH Drive MT Reorganization Similar To DDB. **A)** TIRF-M image of DDH (red) forming clusters (yellow arrows) at MT Minus-ends. Scale bar: 5 μm. **B)** Top: Image frames showing steps during MT reorganization into a mini-aster. Arrows highlight minus-end accumulations that drive sliding in that frame. Bottom: Schematic depicting the DDB-driven movements of MTs. Red dots indicate DDB clusters at minus-ends. Arrows highlight the direction of sliding MTs. Scale bar: 2 μm, time is in sec. **C)** Examples of parallel, anti-parallel, and oblique sliding driven by minus-end accumulations of DDH. Yellow and green arrows indicate minus-ends of sliding microtubules inferred from DDH accumulation. Scale bar: 5 μm, time is in sec. **D)** Examples of mini-asters formed through DDH-driven MT-MT sliding. Scale bars: 5 μm.

**Figure S3:**
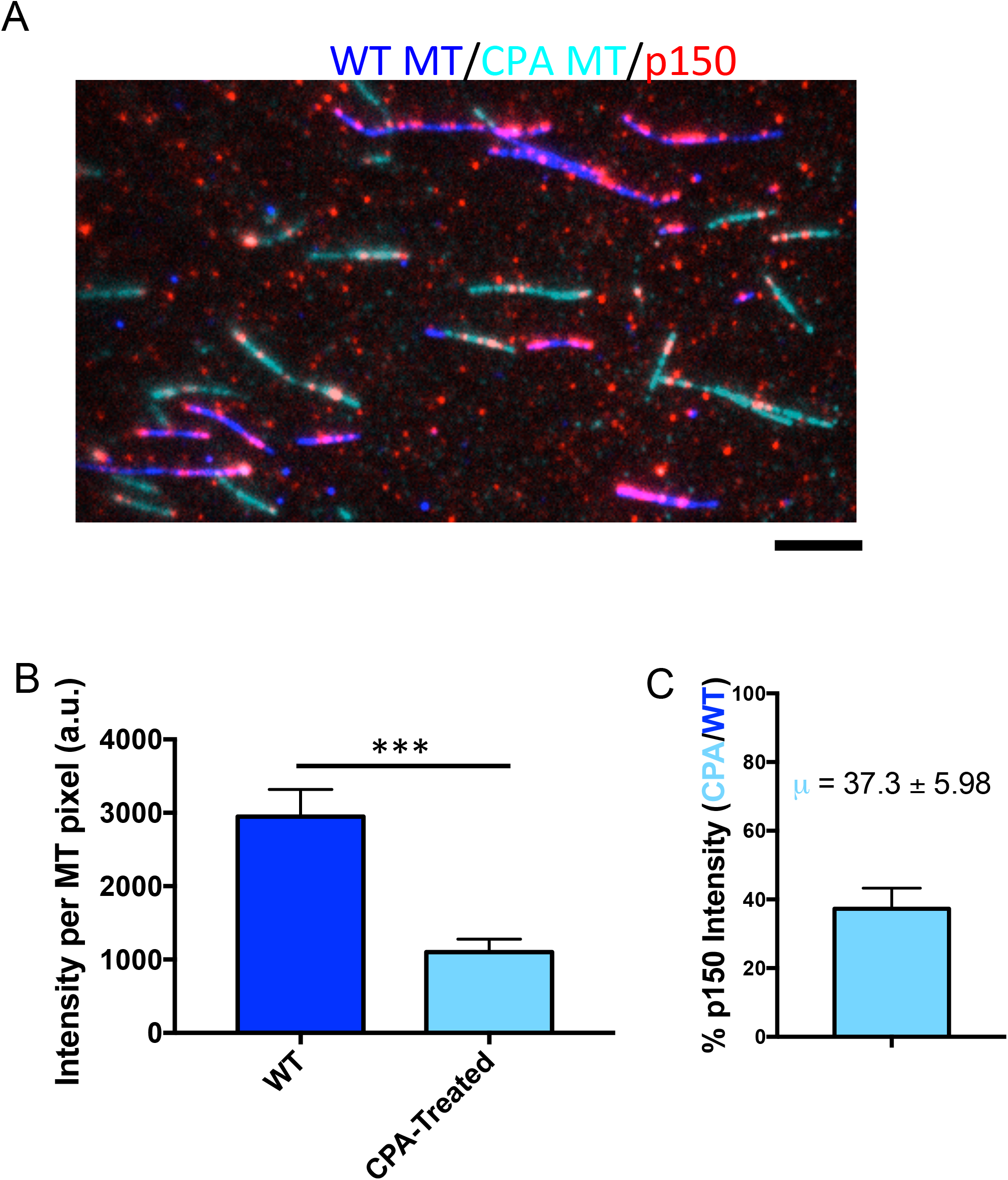
Caboxypeptidase-A treatment of microtubules inhibits binding of recombinant p150^Glued^. **A)** Representative image of TMR-labeled p150-SNAP (red) binding affinity for WT MTs (blue) compared to carboxypeptidase A (CPA)-treated MT (cyan). **B)** Quantification of integrated p150 intensity on either WT or CPA-treated MT. Data represented as intensity minus background per pixel of p150 signal along a microtubule. Average WT and CPA-treated intensity are 2950 a.u. ± 370 and 1100 a.u. ± 176, respectively. P=.0001 by Student’s T-test. Error bars represent SEM. N = 90 and 64 MTs, from two independent experiments. **C)** Quantification of p150 binding to CPA-treated MT versus WT MT. Intensity of CPA-treated microtubules were normalized to the average intensity of p150 binding to WT MT. Data represented as average ± SEM. N = 64.

## Materials and Methods

### Protein Biochemistry

Porcine brain tubulin was isolated using the high-molarity PIPES procedure as described (Castoldi and Popov, 2003) and then labeled with biotin-, Dylight-405 NHS-ester, or Alexa647 NHS-ester as described (http://mitchison.hms.harvard.edu/files/mitchisonlab/files/labeling_tubulin_and_quantifying_labeling_stoichiometry.pdf). Microtubules were prepared by incubation of tubulin with 1mM GTP for 10 min. at 37°C, followed by dilution into 10 μM final taxol for an additional 20 min. For experiments using detyrosinated microtubules, 12 μg/mL carboxypeptidase was added at this step (McKenney et al., 2016). Microtubules were pelleted at 80K rpm in a TLA-100 rotor and the pellet was resuspended in 50 μL BRB80 with 10 μM taxol. StrepII-SNAPf-BicD2N and StepII-SNAPf-Hook3 was isolated from bacteria as described (McKenney et al., 2014). Briefly, a strepII-SNAPf-tagged adaptor constructs were expressed in BL21(DE3) cells (Agilent). The cells were grown at 36° C until OD_600_ of 0.6, then induced with .4 mM IPTG overnight at 18 ° C. Proteins were affinity-purified by streptactin beads, then further purified by size exclusion chromatography on a Superose 6 column in 60 mM Hepes pH 7.4, 50 mM K-acetate, 2 mM MgCl_2_, 1 mM EGTA, 10 % glycerol. Purified BicD2N and Hook3 was used to isolate DDB complexes from rat brain cytosol as previously described (McKenney et al., 2014). DDB complexes were labeled with in a ~4:1 ratio of dye:SNAPf-tagged protein at 2 μM SNAP-TMR, SNAP-Alexa647, or SNAP-Alexa488 dye (NEB) during the isolation procedure, and were frozen in small aliquots and stored at -80°C.

The stoichiometry of labeling and protein concentration was assessed using a Nanodrop One (ThermoFisher) and comparing the absorbance of total protein at 280nm to the absorbance at the SNAP-dye wavelength. Concentrations given are for the total SNAP-labeled amount of protein used in the assay. All buffers and chemicals were from Sigma Aldrich.

### TIRF Microscopy

All microscopy was performed on a custom built through the objective TIRF microscope (Technical Instruments, Burlingame CA) based on a Nikon Ti-E stand, motorized ASI stage, quad-band filter cube (Chroma), Andor laser launch (100 mW 405 nm, 150 mW 488 nm, 100 mW 560 nm, 100 mW 642 nm), EMCCD camera (iXon Ultra 897), and high-speed filter wheel (Finger Lakes Instruments). All imaging was performed using a 100X 1.45NA objective (Nikon) and the 1 or 1.5X tube lens setting on the Ti-E. Experiments were conducted at room temperature. The microscope was controlled with Micro-manager software (Edelstein et al., 2010).

TIRF chambers were assembled from acid washed coverslips (http://labs.bio.unc.edu/Salmon/protocolscoverslippreps.html) and double-sided sticky tape. Taxol-stabilized MTs were assembled with incorporation of ~ 10% Dylight-405-, and biotin-labeled tubulin. Chambers were first incubated with 0.5 mg/mL PLL-PEG-Biotin (Surface Solutions) for 10 min., followed by 0.5 mg/mL streptavidin for 5 min. Unbound streptavidin was washed away with 40 μL of BC buffer (80mM Pipes pH 6.8, 1mM MgCl_2_, 1mM EGTA, 1 mg/mL BSA, 1mg/mL casein, 10μMtaxol). MTs diluted into BC buffer were then incubated in the chamber and allowed to adhere to the streptavidin-coated surface. Unbound MTs were washed away with TIRF buffer (60 mM Hepes pH 7.4, 50 mM K-acetate, 2 mM MgCl_2_, 1 mM EGTA, 10 % glycerol, 0.5 % Pluronic F-127, 0.1 mg/mL Biotin-BSA, 0.2 mg/mL κ-casein, 10μM taxol). Unless otherwise stated, experiments were conducted in imaging buffer (60 mM Hepes pH 7.4, 50 mM K-acetate, 2 mM MgCl_2_, 1 mM EGTA, 10 % glycerol, 0.5 % Pluronic F-127, 0.1 mg/mL Biotin-BSA, 0.2 mg/mL κ-casein, 10μM taxol, 2 mM Trolox, 2 mM protocatechuic acid, ~50 nM protocatechuate-3,4-dioxygenase, and 2 mM ATP).

The resulting data was analyzed manually using kymograph analysis in imageJ (FIJI). For velocity analysis, the velocity of an uninterrupted run segment from a kymograph was used. For images displayed in figures, background was subtracted in FIJI using the ‘subtract background’ function with a rolling ball radius of 50 and brightness and contrast settings were modified linearly. In images where there was substantial drift, the “Descriptor-based series registration (2D/3D + T)” plug-in was used in FIJI with interactive brightness and size detections in the MT channel to stabilize images. Graphs were created using Graphpad Prism 7.0a and statistical tests were performed using this program. All variances given represent standard error of mean.

### Continuous imaging (Drop-in) assay

Accumulation assays were conducted in Nunc Labtek II chambered coverglass system (Thermo Fischer Scientific, #155409). Fixation of microtubules were conducted as before with an adjusted protocol. 50 μL of PLL-PEG was added to the chamber for at least 10 minutes, then aspirated out. Streptavidin, and microtubules were added in the same fashion. Buffer exchanges were done as rapidly as possible to limit surface exposure to air. For imaging, 150 uL of imaging buffer was added in two stages. First, microtubules were aspirated out and 100 uL of imaging buffer was added to the chamber to begin imaging. Next, a 3x dilution of DDB was made in 50 uL of imaging buffer and carefully dropped into the chamber using a pipette during continuous imaging.

### Cluster size quantification

We integrated the fluorescence intensity over a ROI around a MT minus-end cluster. Next 10-30 similar ROIs were drawn around surrounding individual molecules bound to the MT lattice or coverslip to provide an average signal intensity estimate for one DDB molecule. One large ROI was drawn in empty space near the microtubule to provide an accurate estimate for background fluorescence. The average intensity of the background fluorescence ROI was subtracted from the average intensity at minus-ends then integrated over the area of the ROI. This integrated intensity was then compared to the average integrated intensity for one DDB molecule to obtain numbers of DDB at ends.

The kymograph method of determining the number of DDB at minus-ends involved continuous imaging assay as described above. A line scan through the minus-end to obtain an intensity peak over time. The first maximum at saturation was used as a temporal fiducial for when steady state is achieved. We then drew a line scan further into the microtubule lattice and obtained intensity peaks corresponding to DDB traveling across the line to the minus-end. This is the flux of DDB into the minus-end until the time of saturation. This information along with the exponential decay resulting from DDB dwell times in a clustered environment, allowed us to determine the number of molecules at minus-ends at infinity seconds using the formula below.

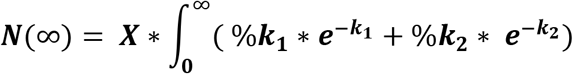

Where N is the number of molecules at minus-ends. X is the rate at which DDB reach minus-ends. The rate constants k_1_ and k_2_ are for short and long dwelling DDB, respectively, and %k_1_ and %k_2_ are the percent of molecules with rate constants k_1_ and k_2_, respectively.

### Single molecule data analysis

Single molecule experiments with DDB and DDH were conducted using chamber slides. DDX at ~1 nM in imaging buffer was flowed into chambers containing fixed taxol-stabilized microtubules. Cluster dwell times were obtained by following the same conditions as above with DDB labeled with either Alexa647- or Alexa488-labled BicD2 at ~30 nM concentrations. Experiments with dynamic microtubules were conducted using coverslip-attached, biotin-labeled GMPCPP stabilized microtubule seeds. Next, imaging buffer containing ~1 nM DDB, 40 μM labeled tubulin (~5% Dylight-405), and 2 mM GTP was introduced into the chamber.

Images were taken 0.5 seconds apart for 1500 frames. Single molecule dwell times were obtained using kymographs. Dwell times were defined as the time when a processive motor became immobile for at the end of a microtubule until it either became diffusive or signal was lost. Processive molecules that immediately dissociated (~1-6% of all molecules) when reaching the end of the microtubule, continued to dwell at the end of the microtubule at the last frame of capture, or overlapped with another molecule were not counted. Images displayed in figures were processed for background subtraction in FIJI using the ‘subtract background’ function with a rolling ball radius of 25-50.

### Sliding Assays and Data Analysis

Experiments were conducted in chamber slides. Fluorescently labeled microtubules diluted in 50 μl of BC+ buffer (BRB80, 1mg/mL casein, 0.2% methylcellulose) was flowed into the chamber. Microtubules not bound to the coverglass were washed out with 20 ul of BC+ buffer. Experiments were conducted in imaging+ buffer (imaging buffer with 0.2% methylcellulose) at 1 second per frame. All angles were defined as the smaller angle between positive ends of microtubules around a central pivot point joining two microtubules. Minus-end foci lifetimes began when minus-ends of two microtubules first contacted each other until they dissociated from one another. Foci that bundled or had another sliding event with a third microtubule were not counted.

### Aster assays

Solutions for aster assays were prepared in 5 μL final volume on ice in TIRF assay buffer with 2 mM ATP, 2 mM GTP, 20 μM Dylight405-labeled tubulin, ~4 μM DDB, and 2 μM taxol. The solution deposited between a slide and coverslip then imaged in widefield at 25° C. DDB were prepared as described previously then concentrated to 10 μM using Amicon spin concentrators with a MW cutoff of 50 KDa (Millipore, UFC510096).

For analysis, background was subtracted in FIJI using the ‘subtract background’ function with a rolling ball radius of 50 and brightness and contrast settings were modified linearly. Standard deviation for the microtubule and DDB channel were obtained by selecting an ROI over the entire field of view. Normalized intensity was normalized to the initial intensity of the first pixel along the line scan.

### Bulk Contraction Assay

Microfluidic devices were prepared as previously described (Foster et al., 2015). Briefly, channel of width 0.9mm and length 18mm were created using AutoCAD 360 (Autodesk) and Silhouette Studio (Silhouette America) software, cut from 125 μm thick tape (3M Scotchcal, St. Paul MN), and adhered to petri dishes. PDMS (Sylgard 184, Dow Corning, Midland, MI) was mixed at a 10:1 ratio, poured onto masters, degassed, and baked overnight at 60° C. Coverslips and PDMS devices were corona treated with air plasma for 1 minute each before bonding. Channels were loaded with a degassed blocking solution composed of 5mg/mL BSA (J.T. Baker, Center Valley, PA) and 2.5% Pluronic-F127 (Sigma, St. Louis, MO) and incubated overnight at 12° C. Channels were further incubated for at least 15 minutes with a solution of 2 mg/mL κ-casein before use.

Taxol stabilized microtubules were made fresh daily as previously described (Nedelec et al., 1997) at a final tubulin concentration of 18.9 μM containing 1.7 μM Alexa-647 or Atto-647 labeled tubulin. Taxol microtubules were diluted 1:19 into a reaction mix such that the final mix was composed of 2.5mM ATP, 20 μM Taxol, DDB at the indicated concentration, and 20% DMSO, all in 1x DDB Buffer. This mix was loaded into a microfluidic channel, sealed using vacuum grease, and imaged immediately using a spinning disk confocal microscope (Nikon Ti2000, Yokugawa CSU-X1), an EMCCD camera (Hamamatsu), and a 2x objective using μManager acquisition software (Edelstein et al., 2010). Images were analyzed using ImageJ and custom written MATLAB software. To extract the contraction timescale and final fraction contracted, ε(t) curves were fit to a saturating exponential function using time points where ε(t) >0.1 as previously described (Foster et al., 2015). All conditions were repeated on at least 3 separate days, using DDB from 2 separate preparations. A similar protocol was used for GST-hDyn experiments, with Atto-488 labeled tubulin substituted as the labeled tubulin and hGST-Dyn1 buffer consisting of 50 mM Tris, pH 8.0 and 150 mM K-acetate substituted for DDB buffer. hGST-Dyn1 experiments were repeated on 3 separate days.

---

**Supplemental Movie S1: DDB dwell and accumulate as stable clusters at microtubule minus-ends**.

**Supplemental Movie S2: Single molecules of DDB dwell at microtubule Minus-ends**.

**Supplemental Movie S3: DDB clusters drive microtubule sliding of all angles**.

**Supplemental Movie S4: DDB clusters drive drastic reorganizations of microtubule bundles**.

**Supplemental Movie S5: DDB accumulate and form MT-MT Foci on growing microtubules**.

**Supplemental Movie S6: DDB create asters from polymerizing MTs**.

**Supplemental Movie S7: Large-scale networks of stabilized microtubules undergo bulk contractions in the presence of DDB but not GST-hDyn**.

**Supplemental Movie S8: DDB localize to the center of MT densities in bulk contractions**.

